# Robust Averaging Protects Decisions from Noise in Neural Computations

**DOI:** 10.1101/147744

**Authors:** Vickie Li, Santiago Herce Castañón, Joshua A Solomon, Hildward Vandormael, Christopher Summerfield

## Abstract

An ideal observer will give equivalent weight to sources of information that are equally reliable. However, when averaging visual information, human observers tend to downweight or discount features that are relatively outlying or deviant (‘robust averaging’). Why humans adopt an integration policy that discards important decision information remains unknown. Here, observers were asked to judge the average tilt in a circular array of high-contrast gratings, relative to an orientation boundary defined by a central reference grating. Observers showed robust averaging of orientation, but the extent to which they did so was a positive predictor of their overall performance. Using computational simulations, we show that although robust averaging is suboptimal for a perfect integrator, it paradoxically enhances performance in the presence of “late” noise, i.e. which corrupts decisions during integration. In other words, robust decision strategies increase the brain’s resilience to noise arising in neural computations during decision-making.

**Author Summary:** Humans often make decisions by averaging information from multiple sources. When all the sources are equally reliable, they should all have equivalent impact (or weight) on the decisions of an “ideal” observer, i.e. one with perfect memory. However, recent experiments have suggested that humans give unequal weight to sources that are deviant or unusual, a phenomenon called “robust averaging”. Here, we use computer simulations to try to understand why humans do this. Our simulations show that under the assumption that information processing is limited by a source of internal uncertainty that we call “late” noise, robust averaging actually leads to improved performance. Using behavioural testing, we replicate the finding of robust averaging in a cohort of healthy humans, and show that those participants that engage in robust averaging perform better on the task. This study thus provides new information about the limitations on human decision-making.

## Introduction

Decisions about the visual world often require observers to integrate information from multiple sources. An ideal observer will give each source a weight that is proportional to its reliability. Thus, where all sources are equally trustworthy, the best policy is simply to average the available features or decision information. For example, a decision about which fruit to buy at the supermarket might involve averaging the estimated size and colour of the produce, or a wager about which football team will win might be made after averaging the speed and skill of all the players on a team [1].

Previous studies have investigated how humans average perceptual information by presenting participants with array composed of multiple visual elements and asking them to report the mean size, colour or shape of the items displayed [2–6]. Interestingly, recent reports suggest that human averaging judgments do not resemble those of an ideal observer [7–10]. Rather, when averaging, humans tend to downweight or discount visual features that are unusual or outlying with respect to the distribution of occurring over recent trials (“robust averaging”). Haberman and Whitney first showed that observers discount emotional deviants when averaging the expression in human faces [7]. Subsequently, de Gardelle, Summerfield and colleagues provided evidence that observers discount outlying colour or shape values during averaging of features in a multi-element array [8, 9]. Control analyses ruled out the possibility that the observed effect was an artefact of hardwired nonlinearities in feature space. Together, these studies suggest that humans are “robust averagers”, overweighting inliers relative to outliers rather than giving equal weight to all elements (although see [11] for a failure to replicate this finding using a 2-alternative forced choice averaging task).

According to a widely-accepted framework with its roots in Bayesian decision theory [1, 12], robust averaging is suboptimal. Intuitively, robust averaging discards information about the stimulus array, and should thus reduce performance relative to a policy that integrates the stimulus feature values evenly. Why, then, do humans give more weight to inliers than outliers during integration of decision information? Here, we tackled this question using psychophysical testing of human observers and computational simulation. We asked participants to average the orientation (tilt) in a circular array of gratings, relative to a central reference grating that either (i) remained the same or (ii) varied in a trial-wise fashion over a block of trials. This latter manipulation allowed us to test whether robust averaging is still observed even when the distribution of sensory information is uniform around the circle and varies randomly from trial. Using this approach, we show that human robust averaging can be conceived of as a policy that rapidly allocates limited resources (gain; see equation 2 below) to items that are closest to the category boundary (or indifference point). Although this policy is suboptimal in the absence of noise, it has a surprising protective effect on decisions that are corrupted by “late” noise arising during or beyond information integration.

Our manuscript is organised as follows. We begin by describing the behaviour of a cohort of human observers performing the orientation averaging task. Next, we describe a simple psychophysical model in which feature values (tilt, relative to a reference value) are transformed nonlinearly before being averaged to form a decision variable. This variable is corrupted with “late” (post-averaging) noise and then used to determine model choices. This model accounts better for human behaviour (including observed robust averaging) than a rival account, based on an ideal observer, that replaces the initial nonlinear step with a purely linear multiplicative transformation. Next, we use simulations to explore the properties of this model. We show that as we increase late noise, a model that engages in robust averaging comes to outperform the linear model, i.e. achieves higher simulated choice accuracy. Finally, we return to the human data, and show that for both model and humans, the use of a robust averaging strategy is a positive predictor of decision accuracy, in particular under high estimated late noise.

## Results

Human participants (*N =* 24) took part in two psychophysical testing sessions separated by approximately one week. On each of 2048 trials, they viewed an array of 8 high-contrast gratings presented in a ring around a single central (reference) grating (**Fig. 1**). The grating orientations were drawn from a single Gaussian distribution with mean μ ∈ {−20°, −10°, 10°, 20°} and standard deviation σ ∈ {8°, 16°} relative to the reference. Their task was to report whether the average orientation in the array was clockwise (CW) or counterclockwise (CCW) of the central grating. The reference grating was drawn uniformly and randomly from around the circle, and varied on either a trial-by-trial (variable reference) or block-by-block (fixed reference) fashion. Fixed and variable reference conditions occurred in different sessions whose order was counterbalanced over participants. Fully informative feedback was administered on every trial.

**Fig. 1.**
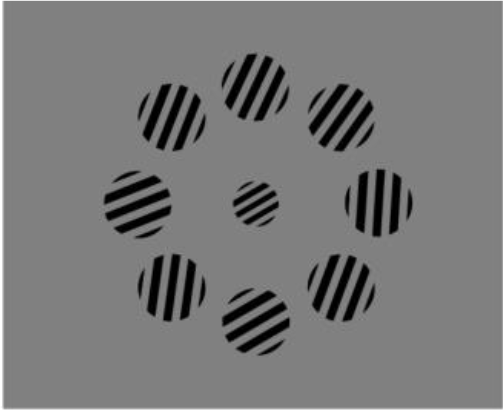
Schematic demonstration of the stimulus array. The task was to report whether the average orientation of the outer ring of gratings fell clockwise or counterclockwise of the orientation of the central (reference) grating.

### Human behaviour

Mean accuracy and standard errors of mean (S.E.M.) for the human participants (lines) are shown in **Fig. 2**. Participants responded more slowly when the orientation mean approached the reference (main effect of |μ|: *F1,20 =* 47.14 *p* < 0.0001) and when the orientation variance increased (main effect of σ: *F1,20 =* 6.84, *p =* 0.017). They also made more errors for lower values of |μ| (F*1,20 =* 397.1, p < 0.0001) and higher values of σ (*F1,20 =* 116.1, *p* < 0.0001). Directly comparing the low |μ| low σ condition (‘low-low’) to the high |μ| high σ condition (‘high-high’), participants made more errors and are slower under high-high condition (accuracy: *F1,20 =* 48.53, p < 0.001; RT: *F1,20 =* 20.67, p < 0.001) even though the|μ| to σ ratio is identical in these two conditions. This result replicates previous findings [8].

**Fig. 2.**
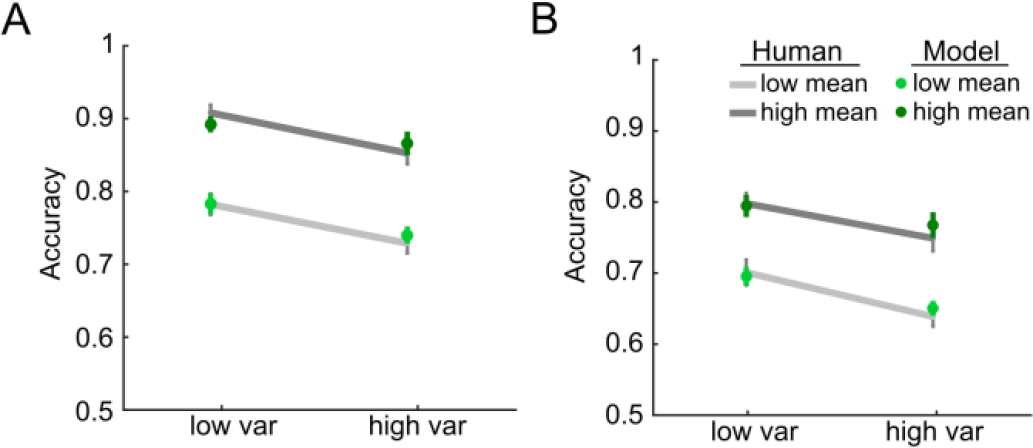
Model and human data. Mean accuracy and the standard error of mean of human (grey lines) and model (green dots) for high and low variance conditions, with low mean (i.e. orientation close to the reference; light grey lines) and high mean (dark grey lines). Panel A shows performance in the fixed reference session, and the panel B shows the variable reference condition.

As expected, participants were overall faster (*F1,20 =* 64.4, *p* < 0.0001) and more accurate (*F1,20 =* 89.95, *p* < 0.0001) in the fixed reference than variable reference condition. An interaction between mean and session was observed for both RT (*F1,20 =* 9.63, *p* < 0.001) and accuracy (*F1,20 =* 5.83, *p =* 0.025) indicated that the cost incurred by lower values of μ was greater under the fixed than variable reference condition. No interactions between session and feature variance were observed. There was a significant interaction for both accuracy (*F1,20 =* 4.18, p *=* 0.41) and RT (*F1,20 =* 8.06, p *=* 0.01) with sessions for the low-low and the high-high condition, showing that the relative performance cost for the high-high condition was lower under the variable reference condition. These findings indicate that our manipulation of fixed vs. variable reference successfully influenced human categorisation performance, and that μ and σ have comparable impact on accuracy and RT to that described in previous studies [8, 9]. The same results were obtained when this analysis was carried out on d’ rather than % correct values (see **Fig. S1**, and **Table S1**).

Next, to probe for robust averaging, we measured the influence that each feature carried on the decision, as a function of its angle relative to the reference (see methods). **Fig. 3A** shows the average regression coefficient (weight) associated with each of 8 bins of the feature values (i.e. orientations relative to reference) for the session with fixed reference (red line) and the session with variable reference (green line). The shaded area shows the standard error of the mean across observers. We first compared the coefficients with a factorial ANOVA, crossing the factors of session (fixed vs. variable reference) and bin. Consistent with the accuracy data above, this yielded a main effect of session (*F1,20 =* 59.54, *p* < 0.001). However, there was also a main effect of bin (*F2.02,40.37=* 6.23, *p =* 0.004) with no interaction between these factors (*p =* 0.31). Next, for each session, we directly compared the weights associated with (i) the four inlying bins (bin 3, 4, 5, 6] and (ii) the four outlying bins (bin 1, 2, 7, 8]. In both sessions, participants gave more weight to those samples falling in inlying than outlying bins (fixed reference: *t20 =* 7.8, *p* < 0.0001; variable reference: *t20 =* 6.3, *p* < 0.0001). In other words, under both fixed and variable reference, participants displayed a pattern of behaviour consistent with a “robust averaging” policy for orientation.

**Fig. 3.**
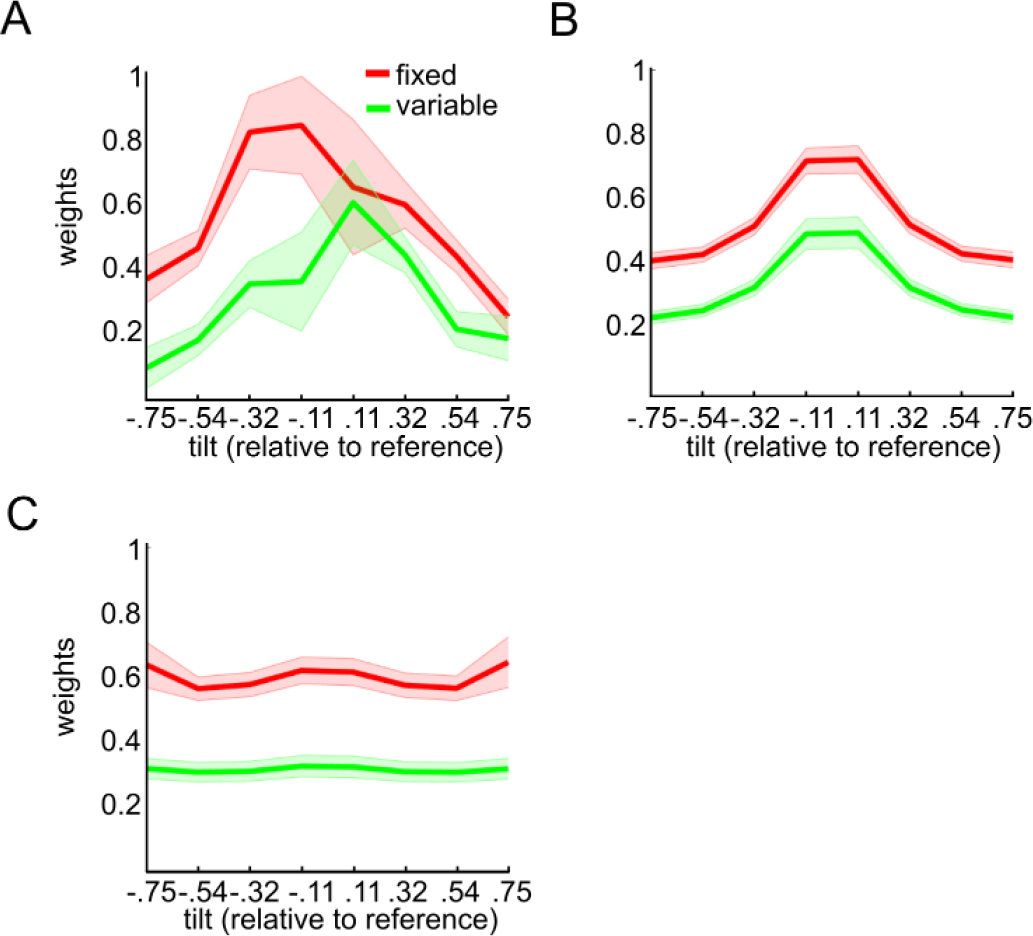
Parameter estimates of orientation of each grating relative to the reference. The y-axis shows parameter estimates for a probit regression in which the angles of orientation of each grating (relative to the reference) were used to predict choice. Angles were tallied into 8 bins, from most negative to most positive relative to the reference, so that each parameter estimate shows the relative weight given to a particular portion of feature space. The x-axis shows the bin center of each bin. The inverted-U shape of the curve is a signature of robust averaging. Shaded areas are the standard error of mean. (A) Weighting functions estimated using human choices (B) Weighting functions for recreated model choices using the best fitting parameters from the power model using the best fitting parameters from human data. (C) Weighting functions for simulated model choice under a case in which angles are linearly mapped onto *DV*

## Model fitting

We fit our data with a simple psychophysical model (power model; see methods). Each array element *i* was characterised by a feature value *X*_*i*_ that was proportional to its orientation, recoded to be relative to the reference (in radians, i.e. in the range −0.79^rad^ to 0.79^rad^ corresponding to −45° to +45°. The model computes a decision value (*DV*) by transforming *X* with a nonlinear function parameterised by an exponent *k*, and summing the resulting values:

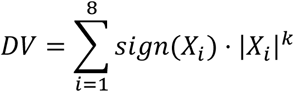

The functions mapping *X* onto *DV* under different levels of *k* (red to blue lines respectively) are shown in **Fig. 4A**. For the special case *k =* 1, the transfer function is linear, and *DV* is equivalent to the simple sum of *X*_*i*_; this is the rule used by the experimenter to determine feedback.

**Fig 4.**
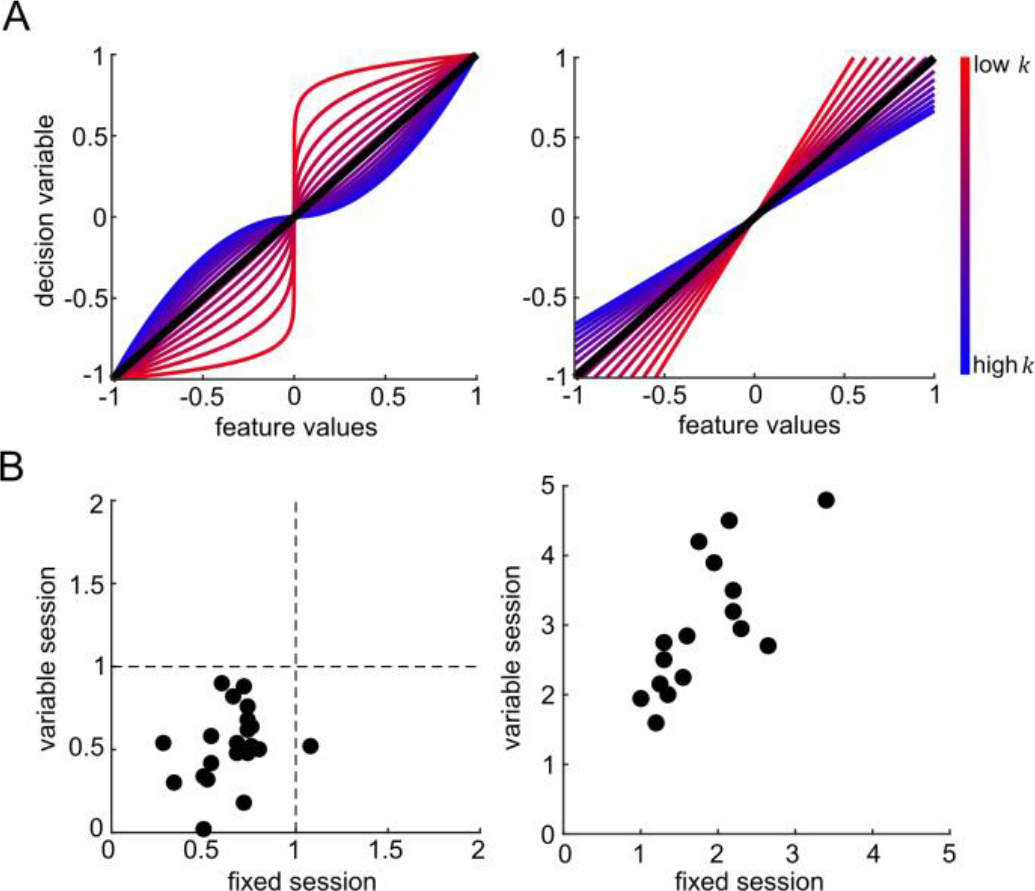

Next, we calculated choice probabilities by passing the *DV* through a sigmoidal choice function with the inverse-slope (*s*; see methods). Varying the inverse slope of the choice function is approximately equivalent to assuming that decision values are corrupted with varying levels of zero-mean Gaussian noise at a post-averaging stage (e.g. “late” noise), with high values of *s* (shallower slope) implying more late noise and thus lower sensitivity. This model allowed us to obtain best-fitting values of *k* and *s* for each participant in both fixed and variable reference conditions, using maximum likelihood estimation. Values of *k* and *s* for each participant are plotted in **Fig. 4B**.

We observed that values for the inverse-slope of the choice function *s* were steeper in the fixed than variable reference condition (*t20 =* 4.27, *p* < 0.001), consistent with lower performance in the variable reference condition. This is likely to reflect the additional processing cost for recoding raw orientations relative to the reference when the latter changed from trial to trial.

Values of *k* did not differ between the fixed and variable reference conditions (*p =* 0.93), but for both conditions, best-fitting values of *k* were lower than 1 (fixed: *t20 =* 9.41, *p* < 0.0001; variable: *t20 =* 3.15, *p =* 0.005). This is consistent with a compression of those array elements that were outlying relative to the reference, i.e. a robust averaging policy. To confirm that the model was showing robust averaging, we then created model choices under the best-fitting parameterisation, by randomly simulating binary choices from the estimates of choice probability using the best-fitting model. Using this approach, we were able to recreate the pattern of accuracy (**Fig. 2**, dots) and weighting profile (**Fig. 3B**) displayed by human participants. In other words, the model displayed comparable costs to humans in each condition, and exhibited the same tendency to engage in robust averaging.

In the model, robust averaging occurs because of the nonlinear form of the function that maps *X*, the feature values, onto *DV*, the decision values, which is steeper in the centre (near 0) and shallower at the edges (far from 0). As a control, we tested the weighting profile observed when *X* is linearly mapped onto *DV*. This confirmed that a linear transformation of feature values did not give rise to robust averaging (**fig. 3C**). Parameter recovery simulation (see methods) confirmed that *k* and *s* were fully identifiable for the power model (shown by **Fig. S2** that actual parameters and recovered parameters fall close to the identity line).

As thus described, our model assumes no noise in the encoding of each individual grating. This assumption follows from the fact that in the experiment, each individual array element (grating) was presented with full contrast and thus the orientation should have been relatively easy to perceive. For example, using a similar stimulus array, one report finds estimates of equivalent encoding noise in the range of 2-6° when contrast values exceed about 0.3 [13]. Moreover, although we additionally randomised the latency with which arrays were presented at 4 levels (250, 500, 750 or 1000 ms). Long presentation latencies led to longer RT on correct choices (*F2.47,56.73 =* 8.65, *p* < 0.001), but this factor had no influence on accuracy (*p =* 0.42; **fig. S3**). Nevertheless, to test this explicitly, we fit a variant of the model in which feature values *X*_*i*_ were corrupted by “early” noise alone – a source of variance that arises before any nonlinearity and averaging, that corrupts each tilt independently relative to the reference (see methods). This model failed to capture the robust averaging effect because the introduction of early noise with power transformation would lead to more stochastic choice pattern. The same feature value that are corrupted by random early noise would drive sometimes the decision to one choice and sometimes to the other choice. We formally compared this “Early noise only” model to our “Late noise only” model, i.e. to that with *k* and *s* described above, finding that it fits the conditionwise accuracy worse in both the fixed reference session (t20 *=* 8.06, p < 0.0001) and the variable reference session (t20 *=* 7.97, p < 0.0001; **Fig. S4C**).

Our model describes the computations that underlie human choices in a simplified fashion, using power-law transducers. However, these functions are intended to describe the output of computations that occur at individual neurons. To demonstrate how transfer functions of this form might arise, we additionally simulated decisions with a population coding model, in which features are processed by a bank of simulated neurons with tuning functions of variable amplitude (see methods). By assuming the height of tuning functions for neurons coding inliers or outliers can vary, we showed in **fig. S5** that we can recreate the family of transfer functions shown in **fig. 4A**. Given that we could recreate the power-law transducer functions using this model, it is unsurprising that the population coding model was also able to recreate the pattern of accuracy (**fig. S6**) and the weighting profile (**fig. S7**) displayed by human participants. However, we chose to model our data with the simpler, psychophysical variant of the model, because it does not require additional assumptions that are not germane to our main points (e.g. the range of tuning widths for the neuronal population).

### Understanding drivers of model performance

Next, turning to our main point, we used simulation to understand how model performance varied under different levels of late noise and degree of robust averaging by exploring different values of *s* and *k*. Model performance (simulated decision accuracy) for the power model under different values of *k* and *s* is shown in **Fig. 5A** (left panel). As expected, performance worsens with increasing late noise (bluish lines). However, performance also depends on *k*. When late noise *s* is higher, the model performs better with lower values of *k* (i.e. those that yield robust averaging). Notably, performance is best with values of *k* that are lower than 1, i.e, under a policy that distorts feature information rather than encoding the feature values linearly.

**Fig. 5.**
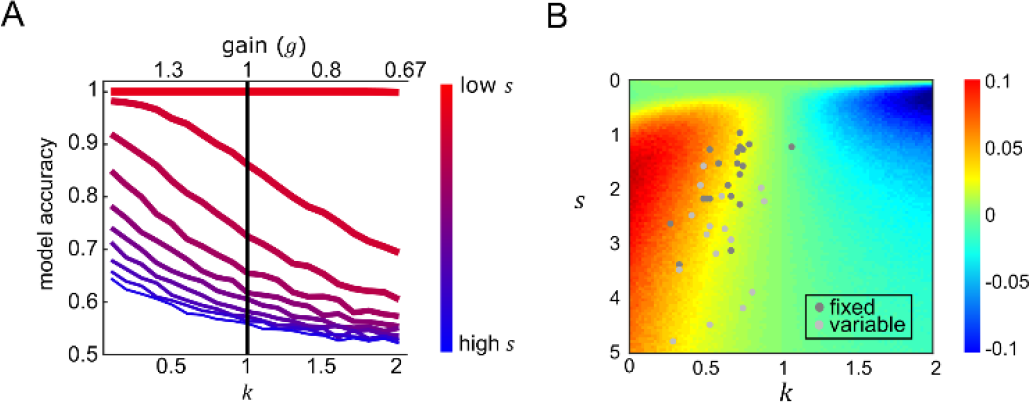
Model accuracy. (A) Simulated model accuracy for the power model under different values of exponent *k* (bottom x-axis, corresponding *g* is plotted on the top x-axis) and late noise (*s*; in a range of 0.05 to 5) in coloured lines with reddish (bluish) lines show simulations with lowest (highest) late noise. The black line is the accuracy of the model when items were allocated with equivalent gain and equally integrated (*k =* 1) (B) After simulating model accuracy of the equivalent gain linear model, performance difference between the power model and the linear model is shown in the coloured surface. Positive values (yellow-red) show parameters where the nonlinear model performance is higher than equivalent linear variants, and negative values (cyan-blue) show the converse. Best fitting *k* and *s* for each subject of the fixed (dark grey dots) and variable reference session (light grey dots) were displayed to show the performance gain relative to using linear weighting scheme.

One trivial reason why model performance might grow as *k* is reduced relates to the scaling of the decision values *DV* that are produced when *X*_*i*_ is transformed. After passage through the sigmoidal choice function, larger values of *DV* will yield choice probabilities that are closer to 0 or 1 and thus increase model performance. To adjust for this, we first calculated the scaling of the decision values that resulted from each transfer function parameterised by a different value of *k*, as follows:

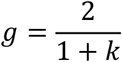

This gain normalisation term is proportional to the integral of the absolute value of the curves in **Fig. 4A**. This normalisation thus adjusts for the expected gain (i.e. proportional increase or decrease in *DV*) that would be incurred by the nonlinear transducer (in the theoretical case in which there is a flat distribution of features). The normalization thus allowed us to compare nonlinear and linear models with equivalent gain. **Fig. S8** shows the resulting value of *g* for each corresponding *k*. We then compared the performance of the model under each transfer function with an equivalent linear model, in which decision values were computed under *k =* 1 (no compression) but rescaled by *g*. This is equivalent to assuming that decisions are limited by a fixed resource (or gain), for example an upper limit on the aggregate firing rates produced by a population of neurons.

Creating this family of yoked linear and nonlinear models allowed us to directly assess the costs and benefits to performance of different values of *k* in a way that controlled for the level of gain. This can be seen in **Fig. 5B**, where we plotted the difference in accuracy between the linear model and a power model that is matched for gain. The red areas in lower left show that when late noise is higher, performance benefits when the model engages more strongly in robust averaging (*k* < 1). In other words, a policy of allocating gain to inliers rather than outliers protects decisions against late noise.

At first glance, this effect might seem counterintuitive. Why should allocating gain preferentially to one portion of feature space prior to averaging benefit performance, if overall gain is equated? One way of thinking about the difference between a power model (with parameter *k*) and a linear model with equivalent gain *g* is that whereas linear model allocates gain evenly across feature space (i.e. equivalently to inliers and outliers), the power model with *k* < 1 focusses gain on those items that are closest to the category boundary, where the transfer function is steepest. Because the overall distribution of features across the experiment is Gaussian with a mode close to the boundary, this means that the power model allocates gain more efficiently, i.e. towards those features that are most likely to occur. We have previously described such “adaptive gain” phenomena in other settings [14, 15].

To verify this contention, we repeated our simulation with a new simulated set of input values *X* that were drawn from a uniform random distribution with respect to the reference, rather than using the Gaussian distributions of tilt values that were viewed by human observers. This simulation revealed no performance advantage for robust averaging. Rather, under uniformly distributed features the best policy was to avoid the nonlinear step and simply average the feature values, as predicted by the ideal observer framework. This is shown in **Fig. 6**, where best performance under the lowest late noise case occurs when feature values are equally integrated. Under high late noise, values of *k* <1 lead to relatively better performance than when all features are equally integrated. However, there is no performance gain for robust averaging compared to the equivalent gain linear model, meaning that unlike in **fig. 5**, the performance gain shown in **fig. 6** is purely due to a larger scaling of input to output values under *k* < 1. This is in fact confirmed by a separate sequential number integration experiment with a different class of stimulus-symbolic numbers. The study showed that that the optimal *k* values under high late noise is greater than 1 since the stimulus were drawn from a uniform distribution [16].

**Fig. 6.**
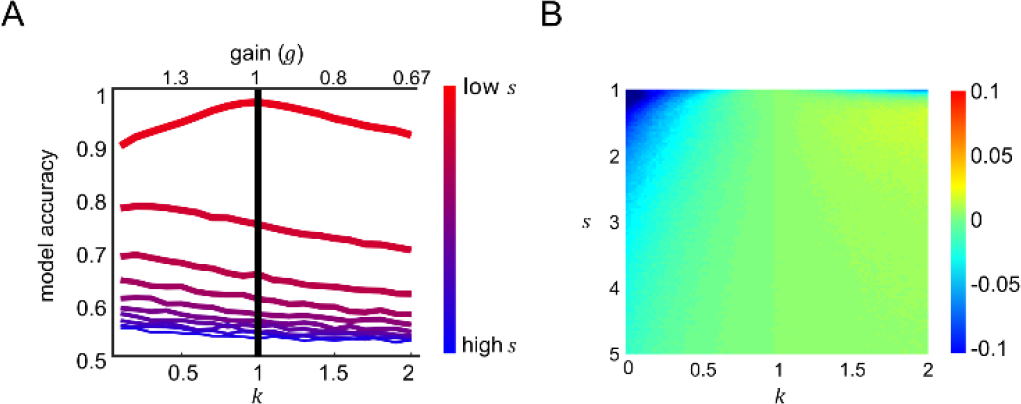
Model accuracy under uniform distributions. Panels A and B are equivalent to panel A and B for Fig 5. However, here the simulations are performed by drawing feature values from uniform random distributions, rather than those used in the human experiment.

#### Linking decision policy to performance

These explorations allow us to make a new and counterintuitive prediction for the human data. If late noise is high, then rather than hurting decision performance, robust averaging should help. We tested this contention using an analysis approach based on multiple regression. For each participant, we split trials into two groups (even and odd). We first obtained the best-fitting *k* and *s* parameters for each participant using even trials. Then, using multiple regression, we estimated multiplicative coefficients that best describe the relationship between the best-fitting parameters for each subject and performance on (left out) odd trials, separately for the fixed and variable reference sessions:

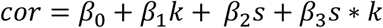

Where *cor* is a vector of mean accuracies (one accuracy for each subject per session), and *k* and *s* are vectors of corresponding best-fitting parameters. In the variable reference condition, both *k* and *s* were significant negative predictors of performance (*k*: *β*_1_ *=* −0.14, *t17 =* −2.51, *p =* 0.022, 95% CI [-0.032 −0.26]; *s*: *β*_2_ *=* −0.041, *t17 =* −7.15, *p* < 0.001, 95% CI [-0.03 −0.052]). In other words, in the variable reference condition, where late noise is intrinsically higher, low values of *k* led to enhanced performance across the human cohort. In the fixed reference session, neither *k* nor *s* was significant predictors of performance (*p =* 0.56 and *p =* 0.16 respectively), but their interaction was significant (*β*_3_ *=* −0.13, *t17 =* −2.88, *p =* 0.01, 95% CI [-0.04 −0.21]). In other words, in the fixed reference condition, predicted performance was higher under lower *k* only for those participants with higher estimated late noise *s*. These findings confirm that in our experiment, robust averaging conferred a benefit on performance under high late noise.

## Discussion

Human observers have previously been shown to be “robust averagers” of low-level visual features such as shape and colour [8, 9], and even of high-dimensional stimuli such as faces [7]. Here, we add to these earlier findings, describing robust averaging of the tilt of a circular array of gratings. However, the focus of the current experiment was to use computational simulations to understand why humans engage in robust averaging. We describe a simple psychophysical model in which features values are transformed nonlinearly prior to averaging. This model assumes the decisions are limited by a fixed resource, and that gain is allocated differentially across feature space, giving priority to inliers – those features that fall close to the category boundary. Through simulations, we find that in our experiment, this relative discounting of outliers gives a boost to performance when decisions are additionally corrupted by “late” noise, i.e. noise arising during, or beyond, the integration of information.

Previously, robust averaging has been considered a suboptimal policy that incurs an unnecessary loss by discarding relevant decision information [17]. The current work offers a new perspective, suggesting that robust averaging is a form of bounded rationality. If we consider an observer whose neural computations are not corrupted by late noise, it is true that robust averaging incurs a cost relative to perfect averaging. However, here we consider decisions as being constrained not just by sources of noise that are external to the observer, or that arise during sensory transduction, but also capacity limits in human information processing. Processing capacity allows a multiplicative gain to be applied to feature values, with higher gain ensuring that feature values are converted to cumulative decision values that fall further from the category boundary (here, the reference orientation). When decision values are further from the category boundary, they are more resilient to “late” noise, which might otherwise drive them to the incorrect side of the category boundary, thereby forcing an error. However, when gain is limited, it must be allocated judiciously. Our simulations show that allocating gain to stimuli that are most likely to occur confers a benefit on performance, and suggest that humans may adopt a robust averaging policy in order to maximise their accuracy on the task.

One longstanding hypothesis states that neural systems will maximise the efficiency of information encoding by allocating the highest resources (e.g. neurons) to those features that are most likely to occur [18]. For example, enhanced human sensitivity to cardinal angles of orientation (those close to 0° and 90°) may reflect the prevalence of contours with this angle in natural scenes [19]. Indeed, neural systems learning via unsupervised methods will naturally learn to represent features in proportion to the frequency with which they occur. Here, we make a related argument for neural gain control. The efficiency of gain control allocation depends on the distribution of features that occurs in the local environment. Allocating gain to features that are rare or unexpected, even when they are more diagnostic of the category, is inefficient, as resources are “wasted” in feature values that are highly unlikely to occur; whereas allocating gain to those features that occur most frequently will confer the greatest benefit. This benefit, however, is only observable when decisions are corrupted by “late” noise, i.e. that arising beyond information averaging. This finding has important implications for our understanding of what may be the “optimal” policy for performing a categorisation task. The ideal observer framework allows us to write down a decision policy that will maximise accuracy for an observer that is limited not by capacity but by noise arising in the external environment. Here, we show an example where the policy that is optimal for an unbiased, noiseless observer is not the one that maximises accuracy for healthy humans.

The current study adds to an emerging body of work that the human brain may have evolved perceptual processing steps that squash, compress or discretise feature information in order to make decisions robust to noise [15]. In another recent line of work, participants were asked to compare the average height of two simultaneously-occurring streams of bars [20] or average value of two streams of numbers [21]. Human choices were best described by a model which discarded information about the locally weaker item, but this “selective integration” policy paradoxically increased simulated performance under higher late noise. As described here, participants seemed to adjust their decision policy to account for their own internal late noise: participants with higher estimated late noise were more likely to engage in robust averaging. Like selective integration, thus, robust averaging is a decision policy that discards decision information but paradoxically confers a benefit on choice.

Additionally, the design of our study allows us to draw conclusions about the timescale over which gain allocation occurs. In previous work, robust averaging was found to vary with the overall distribution of features present in a block of trials. For example, when averaging Gaussian-distributed features in a red-to-purple colour space, purple features were relatively downweighted, but when averaging in a red-to-blue colour space, purple features were relatively upweighted [8]. In other words, the allocation of gain to features depended on the overall distribution of features in the block of trials, with the most frequently-occurring (i.e. expected) items enjoying preferential processing. Here, we saw no difference in robust averaging between a fixed reference condition (in which the Gaussian distribution of orientations remained stable over a prolonged block of trials) and a variable reference condition (in which the Gaussian distribution of orientations changed from trials to trial, and was uniform over the entire session). In other words, any adaptive gain control was set by the reference, and thus occurred very rapidly, i.e. within the timescale of a single trial. Evidence for remarkably rapid adaptive gain control has been described before. Indeed, short-lag repetition priming may be considered a form of gain control [22], in which the prime dictates which features should be processed preferentially [10]. During sequential averaging, the behavioural weight and neural gain applied to a feature depend on its distance from the cumulative average information viewed thus far, as if features pass through an adaptive filter with nonlinear form [14]. These observations are consistent with the theoretical framework that we propose here.

Finally, we discuss some limitations of our approach. Firstly, our model uses a simple power function to describe the nonlinear transformation of inputs prior to averaging. We chose this function for mathematical convenience – it provides a simple means of parameterizing the mapping function feature to decision information in a way that privileges inliers (*k* < 1) or outliers (*k* > 1). However, other forms of nonlinear transformation that are not tested here may also account for the data. Secondly, our best-fitting model assumes zero sensory encoding noise (or ‘early’ noise). Adding early noise to the model did not change qualitatively the benefit of robust averaging under higher late noise, unless it becomes performance-limiting in itself.

However, in other settings, early noise will be an important limiting factor on performance. Although we found that our “late noise only” model fit better than an “early noise only” model, we do not wish to claim that there is no early noise in our task. Since the current experiment was not designed to estimate the level of early noise, it may be of interest to directly manipulate both early and late noise in future experiments.

## Methods

### Ethics statement

The study was approved by the Medical Science Interdivisional Research Ethics Committee (MS IDREC) of the Central University Research Ethics Committee from the University of Oxford. Participants provided written consent before the experiment in accordance with local ethical guidelines.

### Participants

24 healthy human observers (9 males, 15 females; age 23.4±4.7) participated in two testing sessions that occurred one week apart. The order of testing sessions was counterbalanced across participants. The task was performed whilst seated comfortably in front of a computer monitor in a darkened room. Participants received £25 in compensation.

### Task and procedure

All stimuli were created using the Raphaël JavaScript library and presented with the web browser – Chrome Version 49.0.2623.87 on desktop PC computers. The monitor screen refresh rate was 60Hz. Each session consisted of 8 blocks of 128 trials each. On each trial, following a fixation cross of 1000ms duration, participants viewed an array of 8 square-wave gratings with random phase (2.33 cycles/degree, 0.33 RMS contrast, 1.72 degrees visual angle per grating) arranged in a ring 7.82 degrees from the center of the screen (**Fig. 1**). The array was presented for a fixed duration against a grey background in each block (250ms, 500ms, 750ms or 1000ms; this manipulation had little impact on accuracy, and we collapsed across it for all analyses). A single Gabor patch was presented in the centre of the ring contiguous with the array elements (3.49 cycles/degree, 0.33 RMS contrast, 1.15 degrees visual angle). Participants were asked to judge as rapidly and accurately as possible whether the mean orientation of the array of 8 peripheral gratings fell clockwise (CW) or counterclockwise (CCW) of the orientation of the central grating. Feedback was provided immediately following each response: the fixation cross turned green on correct trials for 500ms, and red on incorrect trials for 2500ms. Participants received instructions and completed a training block of 32 trials prior to commencing each session. During the training block, the central grating patch and the array of grating patches remained on the screen for 1 minute or until participants made a response.

### Design

Orientations were sampled from Gaussian distributions with means of *R*+μ where *R* is the reference grating orientation, and variances of σ^2^ on each trial. We crossed μ and σ as orthogonal factors in the design, drawing the orientation mean (in degrees) from μ ∈ {-20°,-10°,10°,20°} and orientation standard deviation σ ∈ {8,16}. Levels of μ and σ are counterbalanced and the order of presentation is randomised across trials in every block. To ensure that the sampled orientations matched the expected distribution with the given μ and σ, resampling of orientation values occurred until the mean and standard deviation of orientation values fell within 1° tolerance of the desired μ and σ. We refer to each of the 8 gratings in the array as a “sample” of feature values. Reference orientations were drawn randomly and uniformly from around the circle. There was a total of 8 blocks per session, leading to a total of 1024 trials per session. In the fixed-reference session, the reference orientation remained fixed over each block of 128 trials. In the variable-reference session, the reference orientation changed from trial to trial. Our experiment thus had a 2 (fixed vs. variable reference) x 2 (μ *=* 10, μ *=* 20) x 2 (σ *=* 8, σ *=* 16) factorial design.

### Analysis

3 subjects were excluded from all analyses due to lowerthan60%accuracy performance in either of the reference condition. Data were analysed using ANOVAs and regressions at the between-subjects (group) level. A threshold of *p* < 0.05 was imposed for all analyses, and we used a Greenhouse-Geisser correction for sphericity where appropriate, so that some degrees of freedom (d.f.) are no longer integers. We first compared accuracy and reaction times for different levels of μ and σ in each session. Next, we used probit regression to estimate the weight with which each sample influenced choices, as a function of its position relative to the reference angle in both fixed and variable reference session. For all analyses, we excluded 13% of trials (‘wraparound’ trials) that contained one or more orientations that were >0.79^rad^ or < 0.79^rad^ (equivalent to >45° or <-45°) relative to the reference, thereby ensuring that we were working within a space in which feature values *X* were approximately linearly related to angle of orientation. A further 0.2% of trials on which no response was registered were also excluded.

For each sample *i* on trial, we assumed that orientations in the sensory space were being recoded as orientations relative to reference in the decision space, and thus refer to the feature values *X* as the orientation relative to the reference. After excluding ‘wraparound’ orientations, all orientations fell within the range of −0.79^rad^ to 0.79^rad^ (equivalent to ±45°). To compute weighting functions, we created for each participant a predictor matrix by tallying values of *X* within each of 8 equally spaced bins (in feature space) with centres between −0.75^rad^ and 0.75^rad^ on a trial-by-trial basis. Values from each bin were entered competitive regressors to regressed against participants’ choices using probit regression. **Fig. 3** is showing the beta weights associated with each bin modulated by the sum of feature values (*X*) within that bin.

### Modelling

#### Power model

Each element *i* was characterised by a feature value *X*_*i*_ in radians (in the range −0.79^rad^ to 0.79^rad^) that was proportional to its orientation relative to the reference. Our model assumes that the decision value (*DV*) that determined choice on each trial was computed by transforming orientations relative to reference using a power-law transducer parameterised by an exponent *k*.

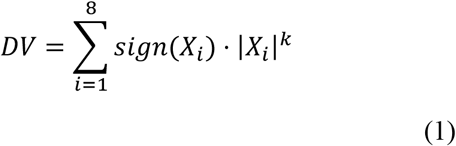

The functions that map feature value *X* onto decision values *DV* for low and high values of *k*. For the special case *k =* 1, the *DV* is equivalent to the simple sum of *Xi*; this is the rule used by the experimenter to determine feedback. Next, we calculated choice probabilities by passing the *DV* through a sigmoidal choice function (see choice probability function and equation 5) with the inverse slope *s*. Higher values of *s* imply shallower slopes and thus greater “late” noise. The sign of sum of *X*_*i*_ always reflect the sign of the mean of the distribution in which *X*_*i*_ was being drawn from, which we used for providing feedback.

#### Equivalent gain factor

Different levels of the exponent *k* vary the convexity or the concavity of the functions shown in Fig. 4a. By considering the integral of the absolute of these functions, it is easy to see that *k* in turn varies the overall scaling of any hypothetically occurring feature values onto *DV*. When *k* < 1, average (absolute) values of *DV* are inflated, and thus pushed away from the category boundary, increasing simulated performance. We wished to ensure that model comparisons cannot be trivially explained by this unequal scaling of feature values to decision variable under different levels of *k*. To correct for this, we thus computed the equivalent gain factor (*g*) that quantifies the average increase in absolute *DV* under different levels of *k*:

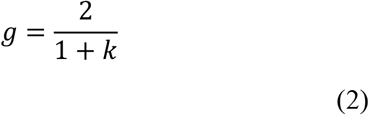

The quantity *g* is equal to 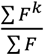 where F is a hypothetical space of features (here, positive only for convenience) that could occur in the experiment. Multiplying equivalent linear models by *g* thus corrects for the inflation that would occur under differing values of *k*. We implemented this correction when comparing equivalent linear and nonlinear models with parameter *k*, either by multiplying the input features of the linear model by *g*, or equivalently, by dividing the output of the nonlinear model by *g*. Importantly, this correction was applied over the features that could occur, not the features that did occur under our mixture of Gaussian-distributed categories. It is for this reason that the nonlinear model leads to improved predicted performance in the experiment we conducted, but not in a simulated experiment in which features were uniformly drawn from across feature space (**Fig. 6**).

#### Equivalent gain linear model

For each nonlinear model variant *k* in the power model, we compute *DV* using a linear model with equivalent gain factor, i.e. a model with the following form:

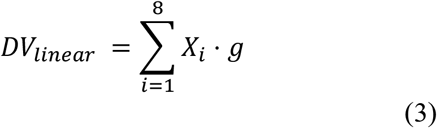

Where *DVlinear* refers to the cumulative decision value of all feature value *Xi* after applied with equivalent gain– *g*. This ensures that each nonlinear power model is compared to a linear model with an equivalent total input-to-output scaling of decision values. Using this approach, we could thus compare the benefits of allocating gain preferentially to inliers (*k* < 1) or outliers (*k* > 1) to allocating gain evenly across feature space (*k =* 1), under the assumption that neural resources were limited to a fixed value defined by *g*, for example the total number of spikes across population of neurons sensitive to orientations. The model comparison of power model against the equivalent gain model is mathematically identical to comparing model performance for *k* < 1 or *k* > 1 against *k =* 1 of a power model which is normalised by *g* in this form:

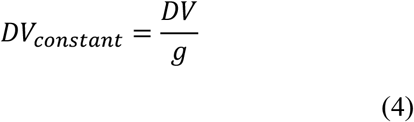

Where *DV constant* refers to the decision variable with probability of a clockwise response (*CW*) given the constant gain across different levels of *k*. Under a *k* < 1 case, inlying items will be allocated with more resources at the expense of depriving resources from outlying items, while under a *k* > 1 case, outlying items will be allocated with more resources at the expense of inlying items. Any difference in simulated model performance of nonlinear transformation of feature values across different values of *k* are not due to differential resources in a linear model.

#### Choice probability function

A choice function with a noise-term *s* was used to transform *DV* of each model into choice probabilities. These choice probabilities are then used for maximum likelihood estimation. We used a choice function of the following form:

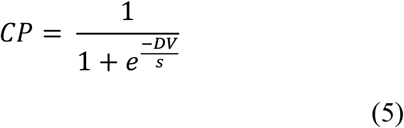

We ensured via visual inspection that the resulting fits were convex over this search space. We then used parametric tests to assess whether the resulting best-fitting parameters differed positively (indicating upweighting of outliers) or negatively (indicating downweighting of outliers) from 1. For each participant, we searched exhaustively over values of *k* (in the range 0.02 to 2) and *s* (in the range 0.05 to 10) that minimised the negative log likelihood of the model.

#### Early noise only model

To test our assumption that early sensory noise (noise arise prior to averaging) alone cannot explain subjects’ choice behaviour, we created a model where each feature value *X*_*i*_ was corrupted by *ε*_*i*_,a sample of noise drawn independently from a Gaussian distribution zero mean and standard deviation *ξ*:

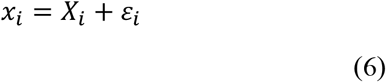

After transforming *x* with exponent *k* using equation 1, we converted the summed of *x* values into a choice probability of 0 or 1 depending of its sign (i.e. via a step function) on a trial-by-trial basis. We fit this model to psychometric functions, by computing the conditional probability of a clockwise response *p(CW)* given the presence of a feature *X*_*i*_ (sorted in to 9 equally spaced bins between −0.75^rad^ to 0.75^rad^). We did this separately for the fixed reference session and variable reference session in humans. Using a grid search method, we identified best-fitting for *ξ* among 20 linearly spaced values from 0 to 3 for each subject and reference condition (fixed, variable) by minimising the MSE between the predicted and observed psychometric functions. **Fig. S4A** shows both human psychometric functions and those predicted by this early noise only model, as well as late noise only model described above, which is parameterised by *k* and *s* (and thus has an equivalent number of free parameters).

Having identified the best-fitting parameters, we used these to predict accuracy for each level of mean and variance, and the weighting function in the fixed and variable reference conditions. The weighting function obtained from best fitting parameterisation of the model is shown on **Fig. S4B** and model fits of accuracies can be seen in **Fig. S4C**. The early noise only model failed to predict the presence of robust averaging and incorrectly predicted that accuracy would not vary as a function of the variance in the stimulus array, and was thus unable to account for human data.

#### Population coding power model

As with the power model, we assume that feature values were recoded from presented orientations relative to the reference into a linear space spanning between –3 and 3 (e.g. radians) where 0 is the value of the reference. We assumed a population of 600 neurons (Μ *=* 600) whose tuning curves are linearly spaced across the feature space. The tuning curve for any neuron, *j*, is defined as a Gaussian probability density function centred at the neuron’s preferred feature value, *f*_*j*_, and with a tuning width fixed across the population, *ε*, specified by an additional free parameter. The amplitude of each neuron’s tuning curve (i.e. its maximum firing rate) was controlled by a gain factor which is a function of the neuron’s preferred feature value, *f*_*j*_, and the power law:

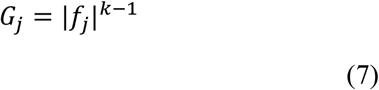

Where *G*_*j*_ represents the gain, *G*, applied to neuron, *j*, whose preferred feature value is *f*_*j*_, and a free parameter, *k*, controls the gain applied across the feature space in the neural population. The firing rate, *R*_*ji*_, for each neuron *j* given a particular stimulus, *X*_i_, is computed as:

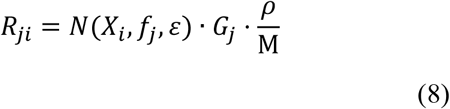

Where (*X*_*i*_, *f*_*j*_, *ε*) correspond to the probability density of a Gaussian with mean, *f*_*j*_, and variance, *ε*, evaluated at point, *X*_*i*_. To adjust for the scaling of output values, the product of the Gaussian density function and gain function is additionally scaled by 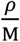, which is the ratio of range of the linear space in radians (*ρ*) to the number of neurons (M). This ensures that the output of the population activity *R* will remained invariant to these factors of no interest in our model. Lastly, the model’s estimate of a stimulus, *X*_*i*_, is a computed from the population of neurons as follows:

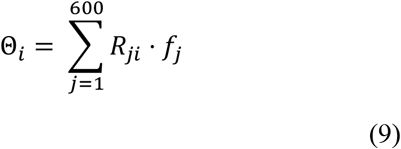

Where *R* is the population activity vector for *X*_*i*_. Firing rate (*R*_*ji*_) of each neuron *j* is weighted by the corresponding neuron’s preferred feature value (*f*_*j*_) before summation to get the model estimate for stimulus (Θ_*i*_). This is then used for computing the cumulative decision values (summation of model estimated angles) on a trial by trial basis for computing choice probability using equation 5 and negative log-likelihood for model fitting.

#### Parameter recovery

To test the ability of the fitting procedure to accurately identify the parameters of the best-fitting power model. We sampled 20 equally-spaced values of *k* (in the range of 0.02 to 2) and *s* (in the range of 0.05 to 10). For each *k* and *s* combination, we transformed a set of orientations presented to subjects in the experiment using the given *k* and computed the choice probability of the *DV* with the given *s*. Then we compared the trial-to-trial estimated choice probability against a random probability drawn from a uniform distribution with a range of 0 to 1 to generate model choices. We then used these artificial choices to recover best-fitting values of *k* and *s* via maximum likelihood estimation.

#### Model performance simulation

We simulated model performance (decision accuracy) under different *k* in a range of 0.02 to 2 and *s* in a range of 0.05 to 5 for the power model. For each combination of *k* and *s*, trial-to-trial estimate of *DV* was computed and transformed into choice probability using equation 5. Model choices were created by comparing the choice probability against a probability drawn randomly from a uniform distribution. Model accuracy was computed as the proportion of model choices that were the same as the pre-defined correct choice, which is simply determined by the sign of the sum of *X*.

## Supporting information

**Fig. S1.**
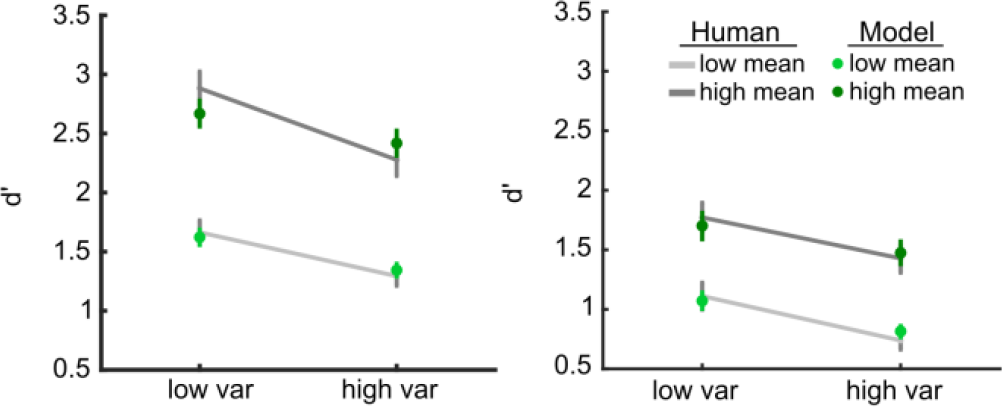
d’ analysis. d’ for each level of |μ| (mean) and σ (variance) conditions were computed separately for fixed reference (Left panel) and variable reference session (Right panel). The grey lines correspond to human’s average d’ for low mean (light grey) and high mean conditions (dark grey). The green dots correspond to the model fits for each condition (low mean in light green dots and high mean in dark green dots).

**S1 table.** ANOVA results on the d’ analysis.

| Variables | d.f. | F values | p-value |
| --- | --- | --- | --- |
| Mean | 20 | 99.37 | $p < 0.001$ |
| Variance | | 292.59 | $p < 0.001$ |
| Session | | 108.41 | $p < 0.001$ |
| Mean x Variance | | 39.2 | $p < 0.001$ |
| Mean x Session | | 4.14 | $p = 0.55$ |
| Variance x Session | | 2.23 | $p = 0.15$ |
| Mean x Variance x Session | | 4.57 | $p = 0.045$ |

**Fig. S2.**
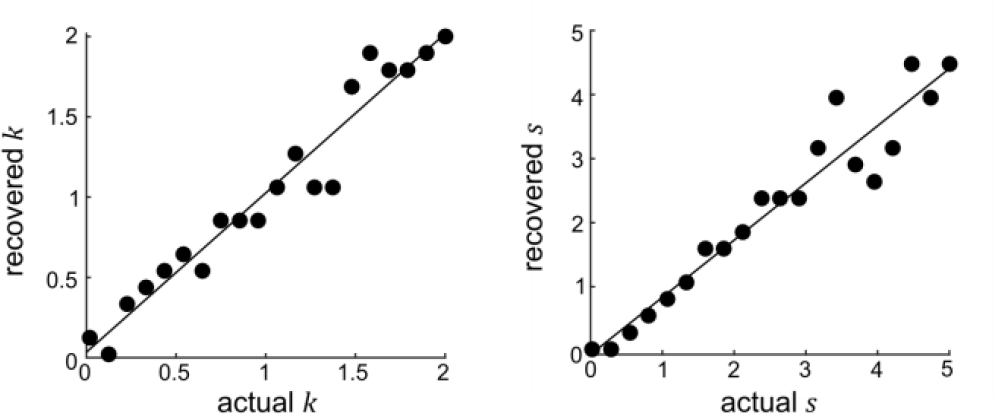
Parameter recovery. Recovered parameters (y-axis) plotted against the actual parameters (x-axis) for *k* (left panel) and *s* (right panel). Black line is the identity line.

**Fig. S3.**
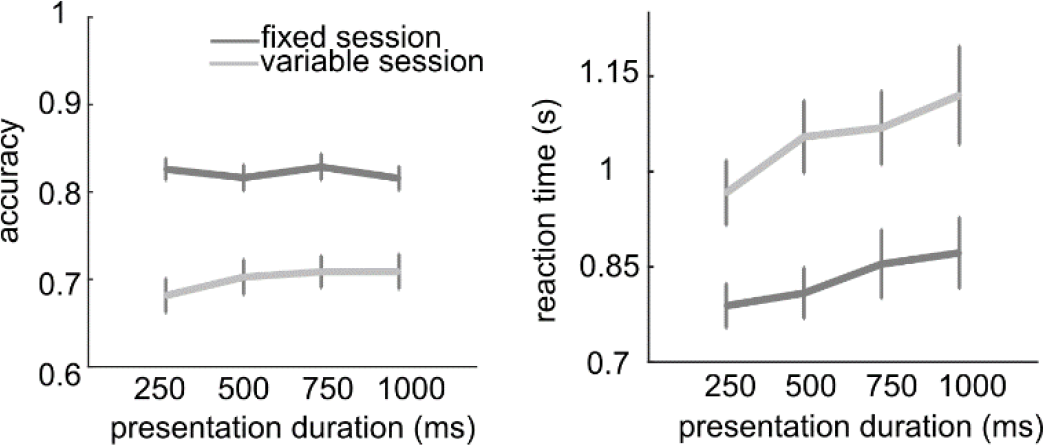
Performance under different presentation duration conditions. Mean and standard error of mean for |μ| on accuracy (left panel) and reaction times (right panel) under different presentation durations (x-axis) in fixed (dark grey line) and variable reference session (light grey line).

**Fig. S4.**
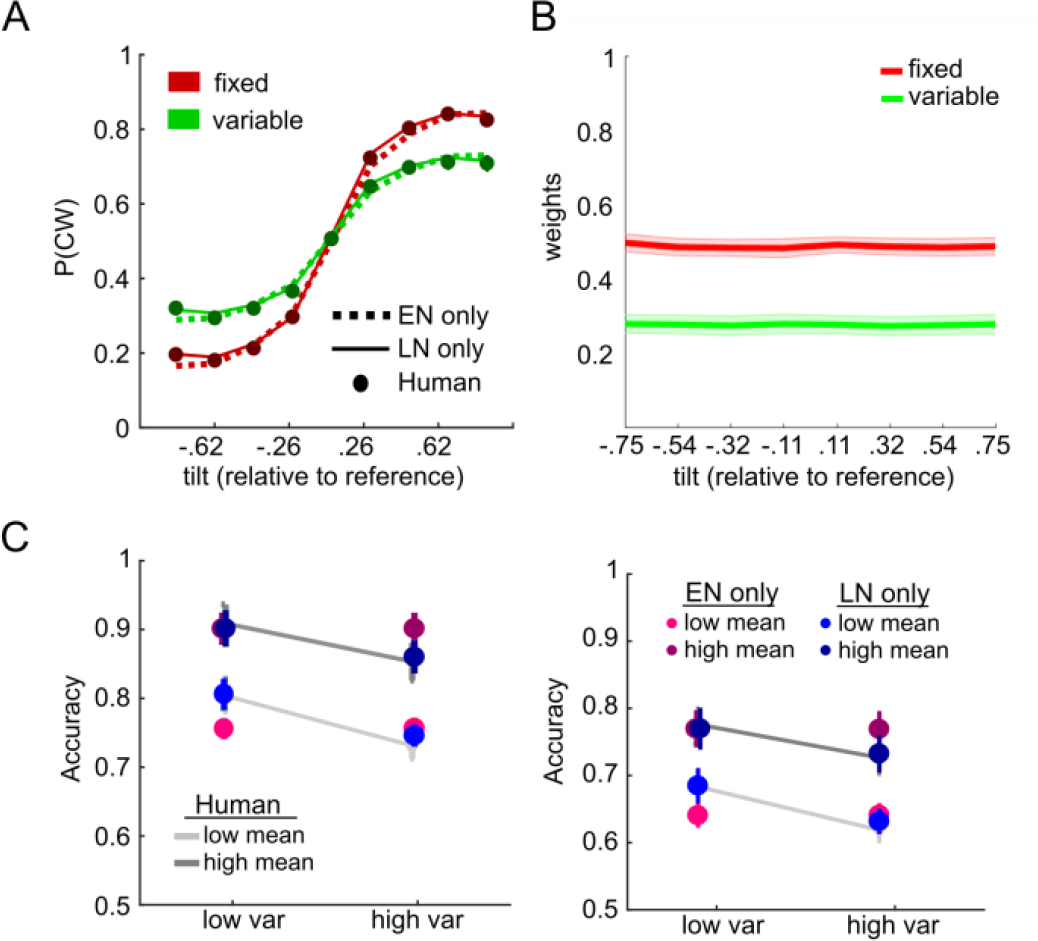
Model comparison of Early noise only model and Late noise only model. (A)Model psychometric functions (dotted line for “EN only” model and thin solid line for “LN only” model) were plotted against humans (darker coloured dots). Both models successfully capture human psychometric functions of the fixed reference and the variable reference sessions (red vs. green). (B) Recreation of the weighting function under simulated choices from the best fitting parameterisation of the early noise model. This model failed to replicate human robust averaging as shown in Fig. 3A. (C) Condition-wise mean accuracy and standard error of mean of the “EN only” model (pinkish dots) and the “LN only” model (bluish dots) superimposed on human accuracies (grey lines). Left panel shows the performance in the fixed reference session, and the right panel shows that of the variable reference condition.

**Fig. S5.**
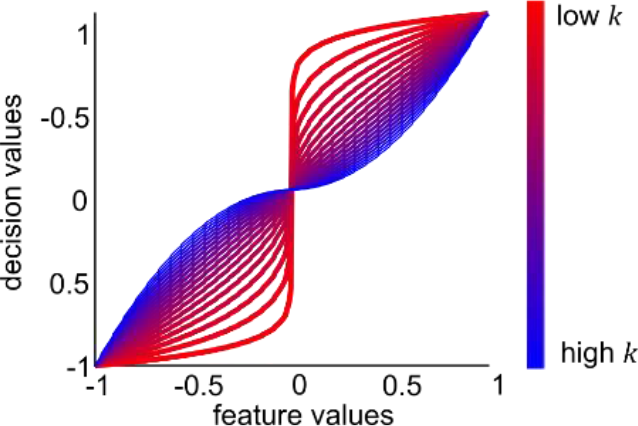
Feature values and decision values generated by a population coding power model. Transfer functions that showed feature values were being transformed into decision values in nonlinear ways under different values of *k* (coloured lines, in a range of 0.02 to 2), similar to transfer functions shown in fig. 4A, which were generated by a simple power model. Tuning width of neurons (*ε*) was assumed to be 0.5 in this illustration.

**Fig. S6.**
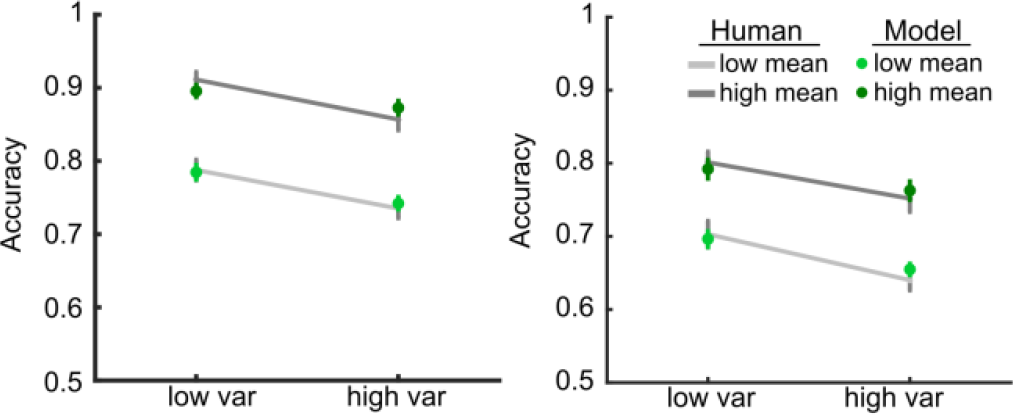
Simulated accuracy under best-fitting parameterisation of population coding. Similar figure shown in fig. 2, this figure is showing the mean (and standard error of mean) accuracy of human (grey lines). Green dots represent the simulated mean accuracy (and standard error of mean) using best-fitting parameters yield from humans with the population coding power model.

**Fig. S7.**
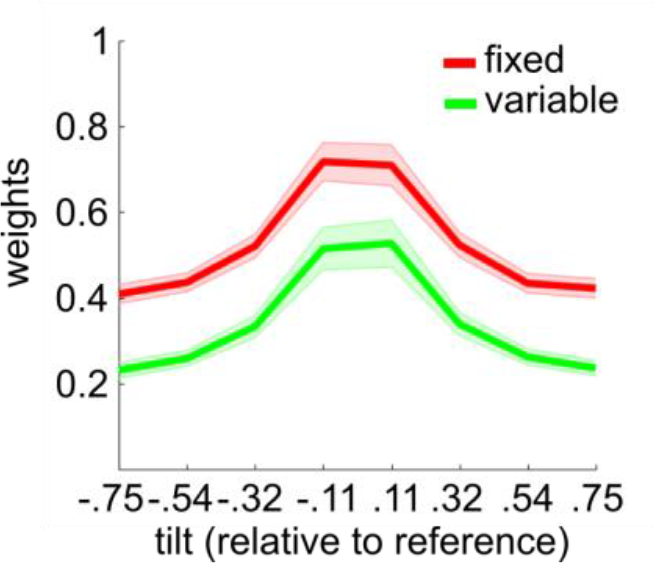
Recreation of parameter estimates using the population coding model. This figure is the same as fig. 3B, but instead of using the simple power model, model choices were simulated using the population coding power model under best-fitting parameterisation of 3 parameters (*ε*, *k*, *s*).

**Fig. S8.**
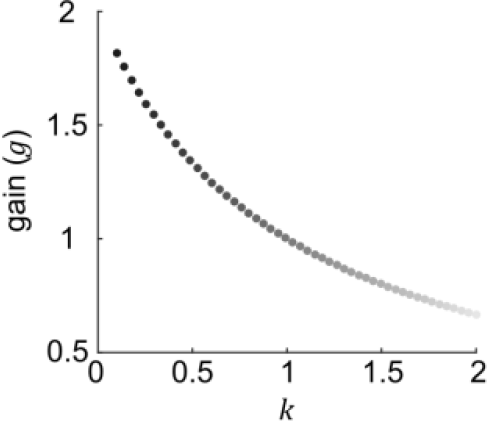
exponent ***k*** and gain (***g***) Lower values *k* (darker dots) have higher multiplicative gain, therefore the corresponding *g* is higher for low value of *k*

## References

1. Wald A, Wolfowitz J. Bayes Solutions of Sequential Decision Problems. Proc Natl Acad Sci U S A. 1949;35(2):99–102. Epub 1949/02/01. PubMed PMID: 16588867; PubMed Central PMCID: PMC1062969.

2. Ariely D. Seeing sets: representation by statistical properties. Psychol Sci. 2001;12(2):157–62. Epub 2001/05/09. PubMed PMID: 11340926.

3. Chong SC, Treisman A. Representation of statistical properties. Vision Res. 2003;43(4):393–404. Epub 2003/01/22. doi: S0042698902005965 [pii]. PubMed PMID: 12535996.

4. Chong SC, Treisman A. Statistical processing: computing the average size in perceptual groups. Vision Res. 2005;45(7):891–900. Epub 2005/01/13. doi: S0042-6989(04)00513-9 [pii] 10.1016/j.visres.2004.10.004. PubMed PMID: 15644229.

5. de Fockert JW, Marchant AP. Attention modulates set representation by statistical properties. Percept Psychophys. 2008;70(5):789–94. Epub 2008/07/11. PubMed PMID: 18613627.

6. Solomon JA, Morgan M, Chubb C. Efficiencies for the statistics of size discrimination. J Vis. 2011;11(12):13. Epub 2011/10/21. doi: 11.12.13 [pii] 10.1167/11.12.13. PubMed PMID: 22011381.

7. Haberman J, Whitney D. The visual system discounts emotional deviants when extracting average expression. Attention, perception & psychophysics. 2010;72(7):1825–38. doi: 10.3758/APP.72.7.1825. PubMed PMID: 20952781; PubMed Central PMCID: PMC3123539.

8. de Gardelle V, Summerfield C. Robust averaging during perceptual judgment. Proc Natl Acad Sci U S A. 2011;108(32):13341–6. Epub 2011/07/27. doi: 1104517108 [pii] 10.1073/pnas.1104517108. PubMed PMID: 21788517; PubMed Central PMCID: PMC3156162.

9. Michael E, de Gardelle V, Nevado-Holgado A, Summerfield C. Unreliable Evidence: 2 Sources of Uncertainty During Perceptual Choice. Cerebral cortex. 2013. doi: 10.1093/cercor/bht287. PubMed PMID: 24122138.

10. Michael E, de Gardelle V, Summerfield C. Priming by the variability of visual information. Proc Natl Acad Sci U S A. 2014;111(21):7873–8. doi: 10.1073/pnas.1308674111. PubMed PMID: 24821803; PubMed Central PMCID: PMC4040545.

11. Gorea A, Belkoura S, Solomon JA. Summary statistics for size over space and time. J Vis. 2014;14(9). doi: 10.1167/14.9.22. PubMed PMID: 25157045.

12. Gold JI, Shadlen MN. Neural computations that underlie decisions about sensory stimuli. Trends Cogn Sci. 2001;5(1):10–6. Epub 2001/02/13. doi: S1364-6613(00)01567-9 [pii]. PubMed PMID: 11164731.

13. Solomon JA, Morgan MJ. Stochastic re-calibration: contextual effects on perceived tilt. Proc Biol Sci. 2006;273(1601):2681–6. doi: 10.1098/rspb.2006.3634. PubMed PMID: 17002955; PubMed Central PMCID: PMCPMC1635463.

14. Cheadle S, Wyart V, Tsetsos K, Myers N, de Gardelle V, Herce Castanon S, et al. Adaptive gain control during human perceptual choice. Neuron. 2014;81(6):1429–41. doi: 10.1016/j.neuron.2014.01.020. PubMed PMID: 24656259.

15. Summerfield C, Tsetsos K. Do humans make good decisions? Trends Cogn Sci. 2015;19(1):27–34. doi: 10.1016/j.tics.2014.11.005. PubMed PMID: 25488076; PubMed Central PMCID: PMC4286584.

16. Spitzer B, Waschke L, Summerfield C. Selective overweighting of larger magnitudes during noisy numerical comparison. Nature Human Behaviour. 2017:Forthcoming.

17. van den Berg R, Ma WJ. Robust averaging during perceptual judgment is not optimal. Proc Natl Acad Sci U S A. 2012;109(13):E736; author reply R7. doi: 10.1073/pnas.1119078109. PubMed PMID: 22362885; PubMed Central PMCID: PMCPMC3324013.

18. Barlow H. Possible principles underlying the transformation of sensory messages. Sensory Communication: MIT Press; 1961.

19. Girshick AR, Landy MS, Simoncelli EP. Cardinal rules: visual orientation perception reflects knowledge of environmental statistics. Nature neuroscience. 2011;14(7):926–32. doi: 10.1038/nn.2831. PubMed PMID: 21642976; PubMed Central PMCID: PMC3125404.

20. Tsetsos K, Moran R, Moreland J, Chater N, Usher M, Summerfield C. Economic irrationality is optimal during noisy decision making. Proc Natl Acad Sci U S A. 2016;113(11):3102–7. doi: 10.1073/pnas.1519157113. PubMed PMID: 26929353; PubMed Central PMCID: PMCPMC4801289.

21. Tsetsos K, Chater N, Usher M. Salience driven value integration explains decision biases and preference reversal. Proc Natl Acad Sci U S A. 2012;109(24):9659–64. doi: 10.1073/pnas.1119569109. PubMed PMID: 22635271; PubMed Central PMCID: PMC3386128.

22. Grill-Spector K, Henson R, Martin A. Repetition and the brain: neural models of stimulus-specific effects. Trends Cogn Sci. 2006;10(1):14–23. Epub 2005/12/03. doi: S1364-6613(05)00323-2 [pii] 10.1016/j.tics.2005.11.006. PubMed PMID: 16321563.

